# An evolutionary analysis of the SARS-CoV-2 genomes from the countries in the same meridian

**DOI:** 10.1101/2020.11.12.380816

**Authors:** Emilio Mastriani, Alexey V. Rakov, Shu-Lin Liu

## Abstract

In the current study we analyzed the genomes of SARS-CoV-2 strains isolated from Italy, Sweden, Congo (countries in the same meridian) and Brazil, as outgroup country. Evolutionary analysis revealed codon 9628 under episodic selective pressure for all four countries, suggesting it as a key site for the virus evolution. Belonging to the P0DTD3 (Y14_SARS2) uncharacterized protein 14, further investigation has been conducted showing the codon mutation as responsible for the helical modification in the secondary structure. According to the predictions done, the codon is placed into the more ordered region of the gene (41-59) and close the area acting as transmembrane (54-67), suggesting its involvement into the attachment phase of the virus. The predicted structures of P0DTD3 mutated and not confirmed the importance of the codon to define the protein structure and the ontological analysis of the protein emphasized that the mutation enhances the binding probability.

## Introduction

The novel coronavirus COVID-19 pandemic caused by SARS-CoV-2 virus currently, in 2020, is the most threatening severe acute respiratory infection in the world. Many attempts to create an effective vaccine against this infection are still in the developing stages and challenging (Bar-Zeev and Inglesby, 2020, Logunov et al., 2020).

The SARS-CoV-2 genome shares about 82% sequence identity with SARS-CoV and MERS-CoV and > 90% sequence identity for essential enzymes and structural proteins. This high level of the sequence identity suggests a common pathogenesis mechanism, thus, therapeutic targeting. Structurally, SARS-CoV-2 contains four structural proteins, that include spike (S), envelope (E), membrane (M), and nucleocapsid (N) proteins (Naqvi et al., 2020). The structure and the genome of the SARS-CoV-2 virus are under extensive study and the results seem to be controversial. It was shown that the two integral membrane proteins (matrix (M) and envelope(E)) tend to evolve slowly by accumulating nucleotide mutations on their corresponding genes, but genes encoding nucleocapsid (N), viral replicase and spike proteins (S), regarded as important targets for the development of vaccines and antiviral drugs, tend to evolve faster (Dilucca et al., 2020). However, other studies showed that potential drug targets of SARS-CoV-2 are highly conserved (Naqvi et al., 2020).

The genome of SARS-CoV-2 is comprised of a single-stranded positive-sense RNA. The newly sequenced genome of the SARS-CoV-2 was submitted in the NCBI genome database (NC_045512.2) ~30 Kb in size. The genetic makeup of SARS-CoV-2 is composed of 13–15 (12 functional) open reading frames (ORFs) containing ~30,000 nucleotides. The genome contains 38% of the GC content and 11 protein-coding genes, with 12 expressed proteins (Naqvi et al., 2020).

The genomic characterization of 95 SARS-CoV-2 genomes revealed three most common mutations that might affect the severity and spread of the SARS-CoV-2(Khailany et al., 2020). Another study highlighted the crucial genomic features that are unique to SARS-CoV-2 and two other deadly coronaviruses, SARS-CoV and MERS-CoV. These features correlate with the high fatality rate of these coronaviruses as well as their ability to switch hosts from animals to humans (Gussow et al., 2020).

The SARS-CoV-2 affected almost every country in the world. COVID-19 has been declared a pandemic and as on 25 October 2020, over 40 million people had been infected and more than 0.934 million deaths.

It is tempting to speculate that the epidemiological and clinical features may differ among different countries or continents.

Genomic comparison of 48,635 SARS-CoV-2 genomes has shown that the average number of mutations per sample was 7.23, and most SARS-CoV-2 strains belong to one of three clades characterized by geographic and genomic specificity: clade G (Europe), clade L (Asia) and G-derived clade (North America) (Mercatelli and Giorgi, 2020).

The obtained results suggest custom-designed antiviral strategies based on the molecular specificities of SARS-CoV-2 in different patients and geographical locations (Mercatelli and Giorgi, 2020). The early study also differentiated three variants according to the geographic location (East Asia, Europe and America) (Forster et al., 2020). More recent genome-wide analysis revealed that frequency of aa mutations were relatively higher in the SARS-CoV-2 genome sequences of Europe (43.07%) followed by Asia (38.09%), and North America (29.64%) while case fatality rates remained higher in the European temperate countries, such as Italy, Spain, Netherlands, France, England and Belgium (Islam et al., 2020).

The aim of study was to compare the set of SARS-CoV-2 genomes from viruses isolated from the countries in the same meridian (longitude): Italy, Sweden, Congo, but with different climate to reveal similar or different pattern of possible evolutionary pressure signatures in their genomes.

## Methods

### Sequence data

We obtained the data from the GISaid repository sampling all the genomes available to that date (May 5, 2020). In details, the file congo-gisaid_hcov-19_2020_05_05_09.fasta with 75 entries, the file italy-gisaid_hcov-19_2020_05_05_10.fasta with 69 entries and the sweden-gisaid_hcov-19_2020_05_05_10.fasta of 104 entries, while the out-group file brazil_gisaid_hcov-19_2020_05_15_04.fasta contained 92 entries. The reference genome with Accession Number NC_045512.2 has been downloaded from the GenBank repository.

### Evolution model analysis

The SARS-CoV-2 Wuhan-Hu-1 genome (RefSeq Acc. No. NC_045512.2) has been used as reference sequence and the VIRULIGN version 1.0.1 application (Libin et al., 2019) to perform the multiple sequence alignment while the AliView version 1.26 application has supported its visualization (Larsson, 2014). HyPhy 2.5.8 (MP) achieved the recombination analysis by the GARD method, the trimming, the removal of the stop codons, the phylogenetic tree and the MEME (Mixed Effects Model of Evolution) analysis (Murrell et al., 2012). The http://vision.hyphy.org/MEME web site has been used to read JSON output files and generate the MEME pictures and tables.

### RNA secondary structure prediction

RNA_fold web server (part of the Vienna RNA Web Suite) predicted the secondary structures of both the sequences (mutated and not) (Lorenz et al., 2011), whereas the graph diagrams have been built using Forna package (Kerpedjiev et al., 2015).

### Protein analysis

The “disorder analysis” of the protein has been conducted by means of MFDp2 (Mizianty et al., 2014), NetSurfP-2.0 (Klausen et al., 2019) and SPOT-Disorder2 (Hanson et al., 2019) applications. The “transmembrane analysis” of the protein has been calculated using the TMHHM server v.2.0, the MemBrain webserver (Yin et al., 2018), the ProtScale (Wilkins et al., 1999) and TMpred (Hofmann, 1993) (scores have been normalized for comparison) on Expasy website (Gasteiger et al., 2003).

### 3D protein structure prediction and ontologies

Both of the protein structures have been calculated with *ab initio* approach using the Robetta webserver (Kim et al., 2004) while DeeProtein capsule from OCEAN CODE (Upmeier zu Belzen et al., 2019) was used to predict the ontologies of the predicted proteins. Three dimensional images of protein structures and their ontologies have been released using PyMOL 2.4.0 (Rigsby and Parker, 2016).

## Results

### Codon 9628 evolves under episodic positive selection

Mixed evolutionary analysis based on MEME algorithm has been conducted on SARS-CoV-2 data from Italy, Sweden, Congo (countries in the same geographic meridian) and Brazil (considered as an outgroup). The investigation revealed the codon 9628 as under episodic positive selective pressure overall the countries, as depicted by the Table 1.

**Table 1.** MEME results. The table reports the data obtained from the evolutionary analysis of SARS-CoV-2 from Brazil, Congo, Italy and Sweden. Here are reported for every country the top 3 sites sorted by p-value. The red color highlights the site 9628.

| ID | site | partition | $\alpha$ | $\beta^-$ | $p^-$ | $\beta^+$ | $p^+$ | LRT | p-value | Branches under selection | Total branch length | MEME LogL | FEL LogL |
| --- | --- | --- | --- | --- | --- | --- | --- | --- | --- | --- | --- | --- | --- |
| BR | 9628 | 1 | 0 | 0 | 0.96 | 10000 | 0.04 | 16.37 | 0 | 2 | 0.65 | -27.28 | -20.62 |
| BR | 9928 | 1 | 0 | 0 | 0.82 | 10000 | 0.18 | 11.12 | 0 | 4 | 2.71 | -31.03 | -28.53 |
| BR | 81 | 1 | 0 | 0 | 0.04 | 1032.18 | 0.96 | 6.95 | 0.01 | 5 | 1.49 | -40.77 | -40.77 |
| CG | 9628 | 1 | 0 | 0 | 0.97 | 10000 | 0.03 | 10.89 | 0 | 1 | 0.25 | -18.18 | -13.54 |
| CG | 2884 | 1 | 0 | 0 | 0.45 | 1273.45 | 0.55 | 3.51 | 0.08 | 5 | 0.60 | -42.49 | -<br>42.37 |
| CG | 6541 | 1 | 0 | 0 | 0.97 | 10000 | 0.03 | 2.73 | 0.12 | 1 | 0.27 | -12.94 | -<br>11.92 |
| IT | 15 | 1 | 0 | 0 | 0.96 | 10000 | 0.04 | 10.21 | 0 | 1 | 0.73 | -15.90 | -<br>12.57 |
| IT | 9628 | 1 | 0 | 0 | 0.97 | 10000 | 0.03 | 11.24 | 0 | 1 | 0.45 | -17.66 | -<br>12.95 |
| IT | 4 | 1 | 0 | 0 | 0.89 | 10000 | 0.11 | 7.25 | 0.01 | 0 | 1.83 | -13.11 | -<br>10.43 |
| SE | 9628 | 1 | 0 | 0 | 0.96 | 9613.52 | 0.04 | 16.03 | 0 | 2 | 0.51 | -27.43 | -<br>21.10 |
| SE | 4409 | 1 | 0 | 0 | 0.97 | 4356.70 | 0.03 | 7.68 | 0.01 | 1 | 0.16 | -15.63 | -<br>12.33 |
| SE | 4732 | 1 | 0 | 0 | 0.95 | 10000 | 0.05 | 3.85 | 0.07 | 2 | 0.74 | -19.66 | -<br>18.78 |

In this context we use the term “site” as synonymous of codon, respecting the HyPhy terminology. The asymptotic p-value for episodic diversification at site 9628 results equal to zero in all cases and Figure 1 shows the distribution of the p-value over the sites for all countries.

**Figure 1.**
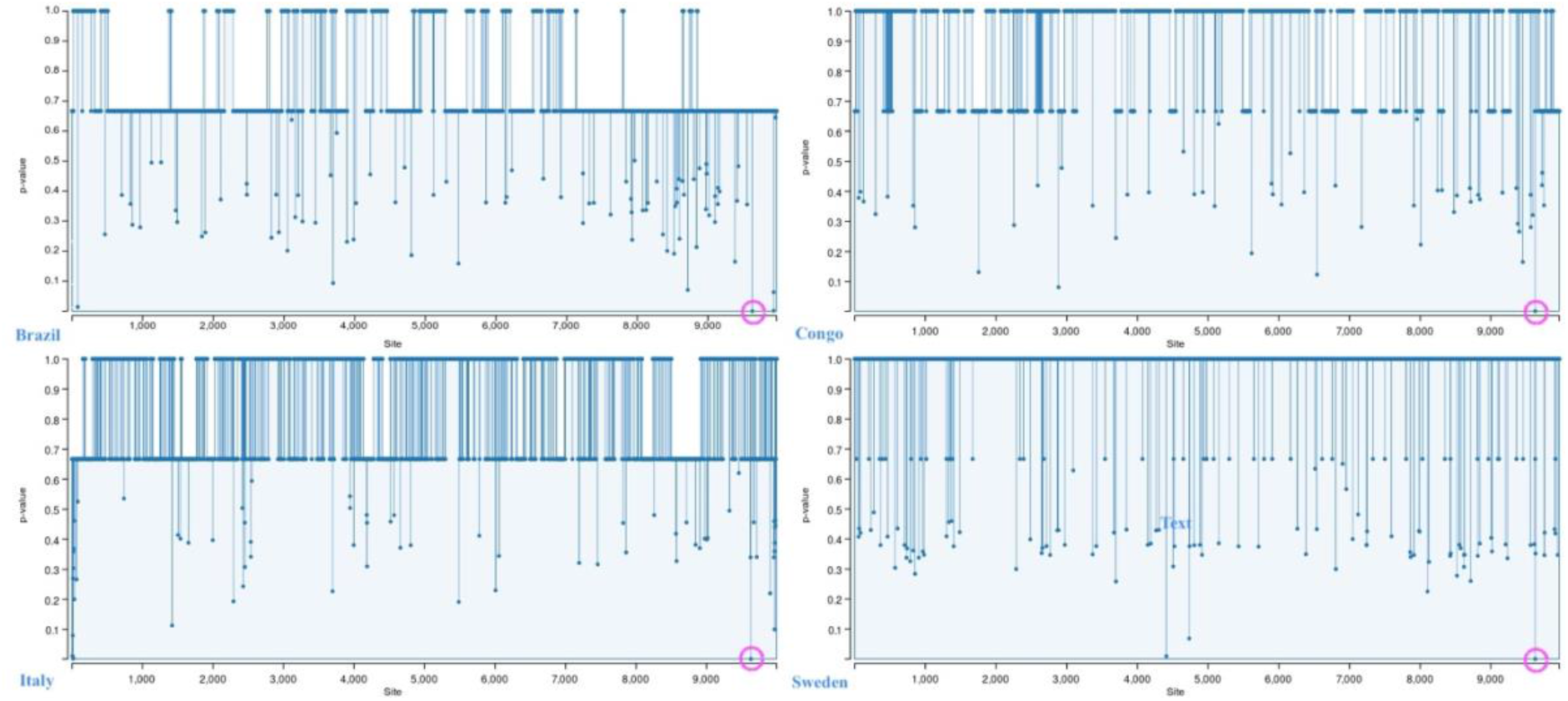
MEME site plot. Distribution of the p-value over the sites of the Brazil, Congo, Italy and Sweden countries. The purple circle indicates the 9628 site under episodic selective pressure.

The deep check of the multiple alignment data of the countries revealed the episodic positive selective pressure on site 9628 as a consistent mutation of the codon GGG to ACG, as showed in the Figure 2.

**Figure 2.**
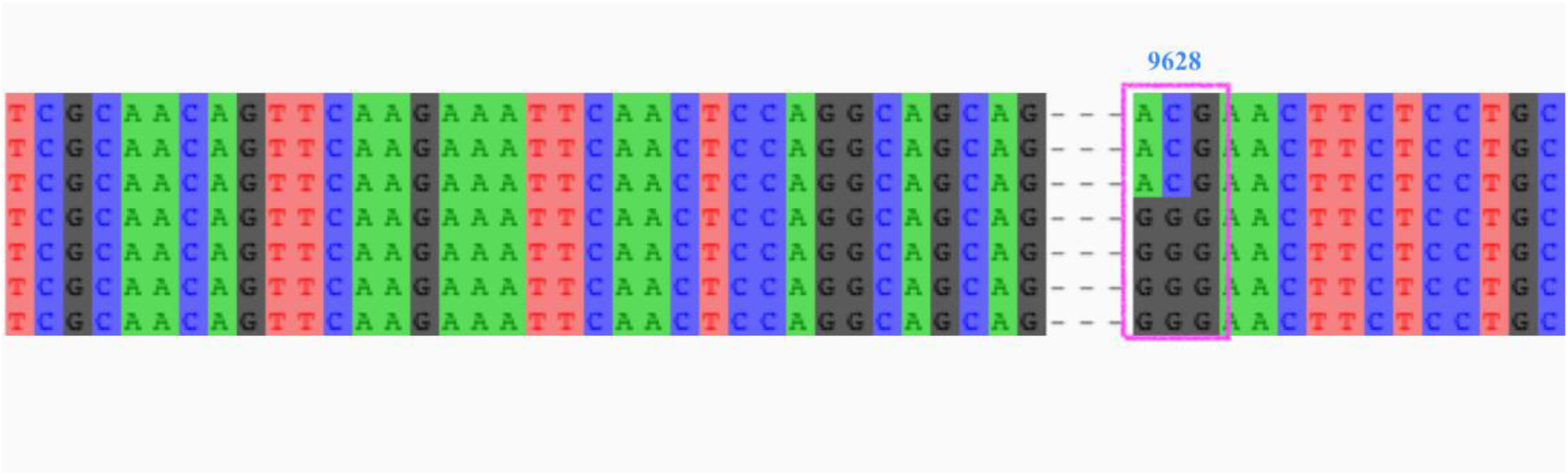
Site 9628 on MSA. Part of the multiple sequence alignment from the Italian data showing the site 9628 under episodic selective pressure. The nucleotides mute from GGG to ACG

### RNA secondary structure prediction changes

The secondary structure prediction before and after mutation shows important differences. In detail, the mutation from GGG to ACG has been applied to the RNA sequence as shown in the Figure 3. The comparison between the prediction of the two secondary structures highlighted structural modification at the top-right ring of the RNA-conformation, as depicted in the Figure 4. This result induces to suppose the GGG to ACG mutation as responsible for a significant modification of the RNA secondary structure.

**Figure 3.**
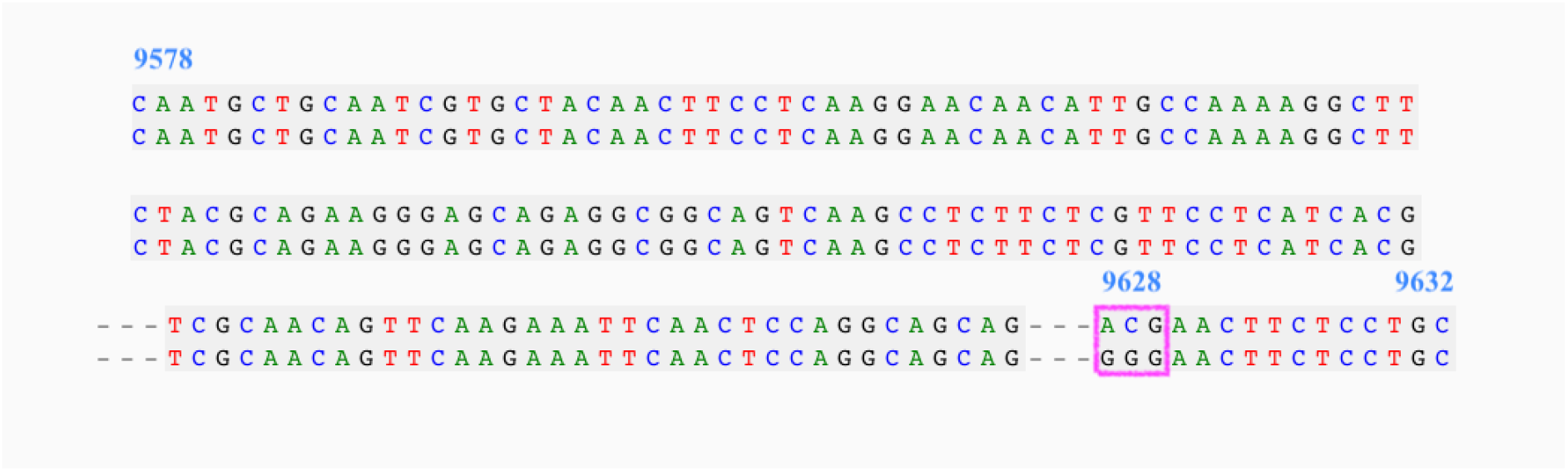
Nucleotides mutation over aligned sequences. The sequence considered to predict the secondary structures in case of mutation and not. Site position is typed in blue, from the start codon (9578) to the ORF (9632)

**Figure 4.**
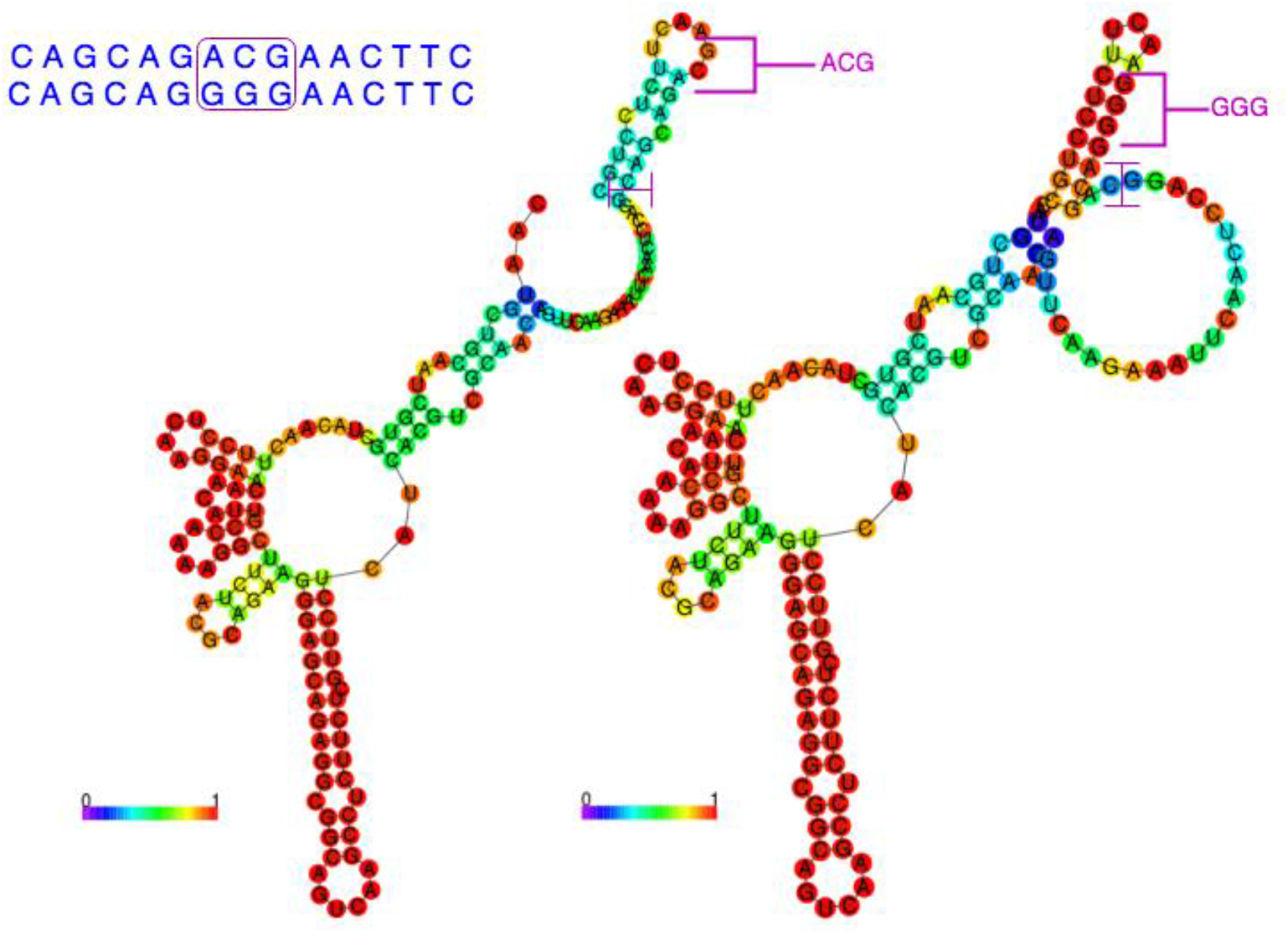
Secondary structure prediction. The two RNA-diagrams exhibit the structural modification affected by the GGG to ACG mutation.

### Protein analysis

The analysis of the protein conducted for finding its disordered region, turns out the positions from 41 to 59 as the more stable, with the Glycine (G) amino-acid placed at the 50th. We gained the results by using three different software and considering the average value for the probability of disorder, as reported by the Table 2 and represented in the Figure 5. Further analysis to locate the transmembrane region in the protein, revealed the locations from 54 to 67 as associated to this function. The analysis, conducted with the utilization of four distinct web applications and by evaluating their average values, puts the Glycine (G) amino-acid as fairly near to the transmembrane region to suppose its involvement. Table 3 reports some of the data showing the probabilities of acting as transmembrane for each aminoacidic position, while Figure 6 illustrates the distribution of the probabilities by remarking the region mentioned above.

**Figure 5.**
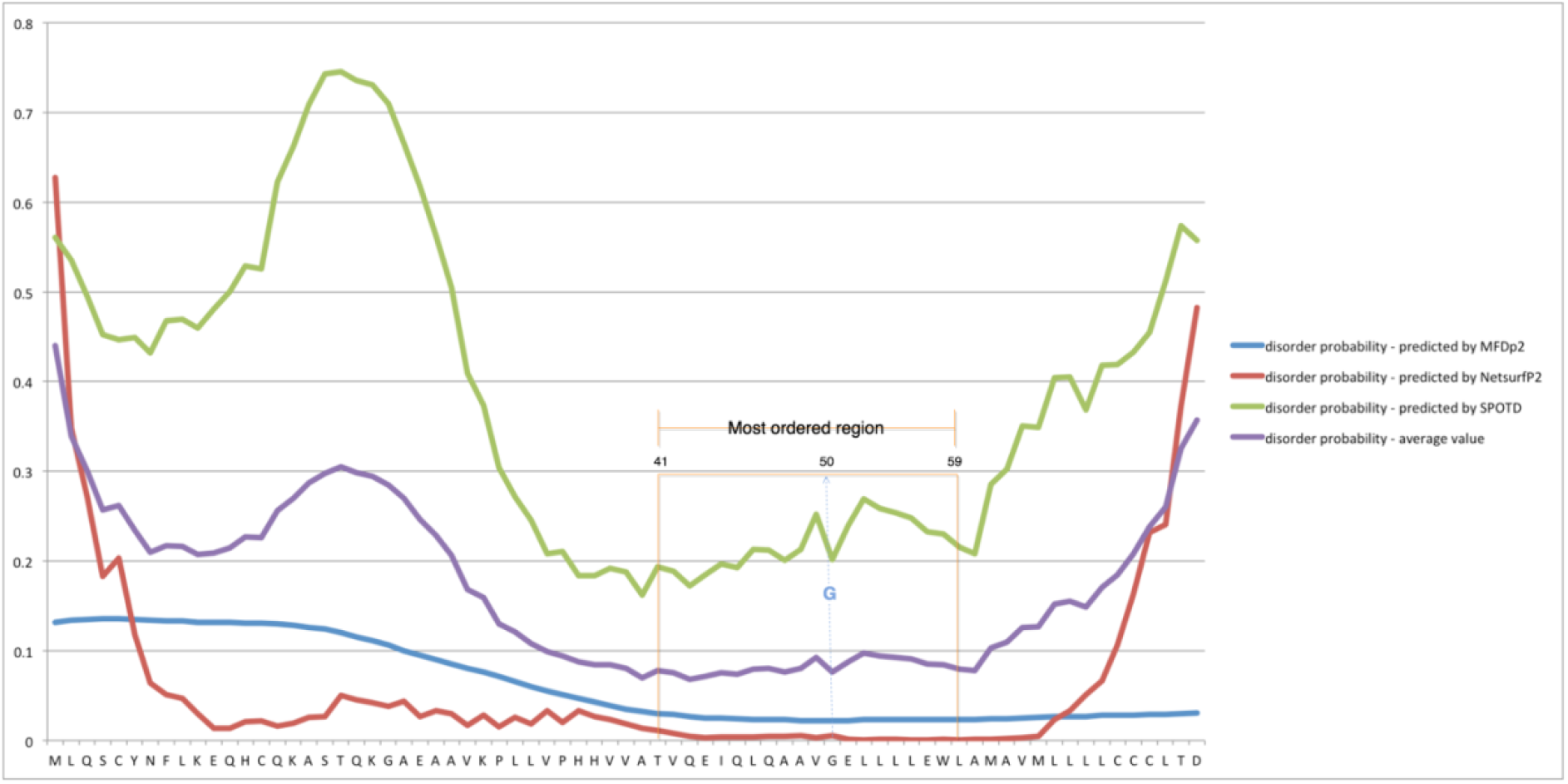
Disorder region analysis. The region 41-59 results as that with lowest probability to be disordered. The orange rectangle delimits the region and the blue dots outline the position of G on the different curves

**Figure 6.**
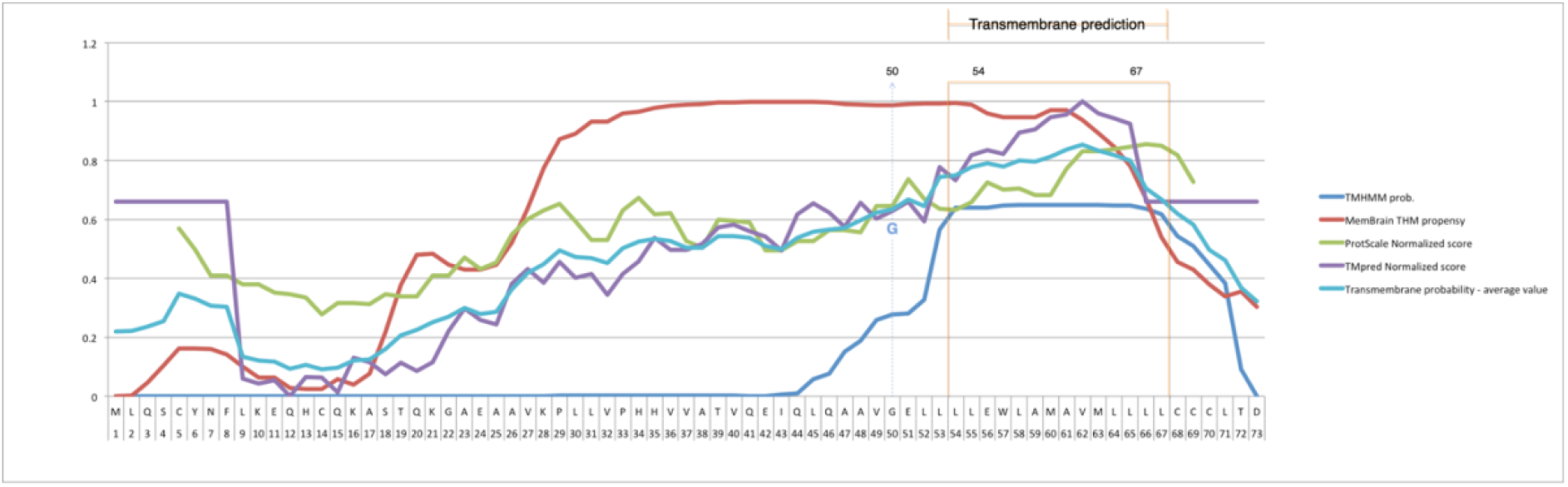
Transmembrane prediction. The region 54-67 results as that one with highest probability to code for the transmembrane and G amino-acid is near enough to suppose its involvement. The orange rectangle delimits the region and the blue dots outline the position of G on the different curves

**Table 2.** Disorder analysis. The table reports the probability of disorder for each position of the protein. The probabilities have been calculated using MFDp2, Netsurf and SPOTD software. The last column assays the average value of the disorder probability for each position. In red the amino-acid G placed at position 50, inside of the stable region.

| *Position* | **AA**<br>**Sequence** | **Dis. Prob.**<br>**MFDp2** | **Dis. Prob.**<br>**NetsurfP2** | **Dis. Prob.**<br>**SPOTD** | **Dis. Prob. average**<br>**value** |
| --- | --- | --- | --- | --- | --- |
| 1 | M | 0.132 | 0.627823114 | 0.5607 | 0.440174371 |
| 2 | L | 0.134 | 0.347978383 | 0.5358 | 0.339259461 |
| 3 | Q | 0.135 | 0.270706475 | 0.4945 | 0.300068825 |
| ... |  |  |  |  |  |
| 39 | T | 0.03 | 0.010842944 | 0.1936 | 0.078147648 |
| 40 | V | 0.029 | 0.007660664 | 0.189 | 0.075220221 |
| 41 | Q | 0.027 | 0.004478907 | 0.172 | 0.067826302 |
| 42 | E | 0.025 | 0.00340931 | 0.1848 | 0.07106977 |
| 43 | I | 0.025 | 0.003887762 | 0.1968 | 0.075229254 |
| 44 | Q | 0.024 | 0.003997837 | 0.1927 | 0.073565946 |
| 45 | L | 0.023 | 0.00361518 | 0.2129 | 0.079838393 |
| 46 | Q | 0.023 | 0.004551574 | 0.2123 | 0.079950525 |
| 47 | A | 0.023 | 0.004939525 | 0.2011 | 0.076346508 |
| 48 | A | 0.022 | 0.005752307 | 0.2133 | 0.080350769 |
| 49 | V | 0.022 | 0.002826149 | 0.2524 | 0.092408716 |
| 50 | G | 0.022 | 0.005828088 | 0.2013 | 0.076376029 |
| 51 | E | 0.022 | 0.001046103 | 0.24 | 0.087682034 |
| 52 | L | 0.023 | 0.000922468 | 0.2694 | 0.097774156 |
| 53 | L | 0.023 | 0.001263275 | 0.2588 | 0.094354425 |
| 54 | L | 0.023 | 0.001187441 | 0.2539 | 0.092695814 |
| 55 | L | 0.023 | 0.000650476 | 0.2483 | 0.090650159 |
| 56 | E | 0.023 | 0.000615434 | 0.2328 | 0.085471811 |
| 57 | W | 0.023 | 0.001080571 | 0.2302 | 0.08476019 |
| 58 | L | 0.023 | 0.000941573 | 0.2154 | 0.079780524 |
| 59 | A | 0.023 | 0.001573079 | 0.208 | 0.07752436 |
| 60 | M | 0.024 | 0.000997698 | 0.2853 | 0.103432566 |
| 61 | A | 0.024 | 0.00227783 | 0.3026 | 0.109625943 |
| 62 | V | 0.025 | 0.003362786 | 0.3503 | 0.126220929 |

**Table 3.** Transmembrane prediction. Results have been obtained using TMHMM, MemBrainTHM, ProtScale and TMpred applications. Results from ProtScale and TMpred have been normalized for comparison with other probabilities. The last column reports the average value of probability for each position.

| Position | AA<br>Sequence | TMHMM<br>M prob. | MemBrain<br>THM<br>propensity | ProtScale<br>Normalized<br>score | TMpred<br>Normalized<br>score | Transmembrane<br>probability - average<br>value |
| --- | --- | --- | --- | --- | --- | --- |
| 1 | M | 0 | 0.000191 |  | 0.661425764 | 0.220538921 |
| 2 | L | 0 | 0.002851 |  | 0.661425764 | 0.221425588 |
| 3 | Q | 0 | 0.046538 |  | 0.661425764 | 0.235987921 |
| ... |  |  |  |  |  |  |
| 49 | V | 0.2594 | 0.987914 | 0.646 | 0.603358942 | 0.624168236 |
| 50 | G | 0.27719 | 0.987914 | 0.646 | 0.629801679 | 0.63522642 |
| 51 | E | 0.28083 | 0.991702 | 0.736 | 0.660532428 | 0.667266107 |
| 52 | L | 0.32735 | 0.993857 | 0.67 | 0.594246918 | 0.646363479 |
| 53 | L | 0.56651 | 0.993857 | 0.637 | 0.778452743 | 0.743954936 |
| 54 | L | 0.63937 | 0.994522 | 0.632 | 0.73360729 | 0.749874822 |
| 55 | L | 0.64032 | 0.990459 | 0.659 | 0.818831517 | 0.777152629 |
| 56 | E | 0.64052 | 0.96027 | 0.726 | 0.835626228 | 0.790604057 |
| 57 | W | 0.64826 | 0.946819 | 0.701 | 0.822583527 | 0.779665632 |
| 58 | L | 0.6493 | 0.947424 | 0.706 | 0.895122387 | 0.799461597 |
| 59 | A | 0.64928 | 0.947424 | 0.683 | 0.905663748 | 0.796341937 |
| 60 | M | 0.64927 | 0.970735 | 0.683 | 0.947293193 | 0.812574548 |
| 61 | A | 0.64924 | 0.970735 | 0.773 | 0.955511881 | 0.83712172 |
| 62 | V | 0.64903 | 0.937507 | 0.831 | 1 | 0.85438425 |
| 63 | M | 0.64893 | 0.892506 | 0.831 | 0.960871896 | 0.833326974 |
| 64 | L | 0.6482 | 0.846403 | 0.84 | 0.942826514 | 0.819357379 |
| 65 | L | 0.64758 | 0.781733 | 0.847 | 0.924066464 | 0.800094866 |
| 66 | L | 0.63557 | 0.670387 | 0.856 | 0.661425764 | 0.705845691 |
| 67 | L | 0.61835 | 0.539353 | 0.851 | 0.661425764 | 0.667532191 |
| 68 | C | 0.5428 | 0.455615 | 0.819 | 0.661425764 | 0.619710191 |
| 69 | C | 0.51009 | 0.430385 | 0.728 | 0.661425764 | 0.582475191 |
| 70 | C | 0.44702 | 0.380525 |  | 0.661425764 | 0.496323588 |

### 3-Dimensional Protein analysis

Being P0DTD3.1 an uncharacterized protein, it has been necessary to predict the three-dimensional structure using the *ab initio* approach for both the protein sequences, mutated and not. According to the preliminary clue from the secondary structure prediction, the model of the mutated protein presents a slightly different structure around the aminoacidic residue changing from G to T. The Figures 7 and 8 illustrate the predicted two models showing as the mentioned mutation impacts the tertiary structure of the protein. In fact, the comparison between the residues 45-55 in MUT31136 and MOD30336 turns out that portion of the protein subject to the mutation stretches out with repercussions to the preceding helix. This result suggests that the mutation of the single amino-acid from G to T with the consecutive stretching upon the 3D structure of the protein tends to make the protein acquire new functions.

**Figure 7.**
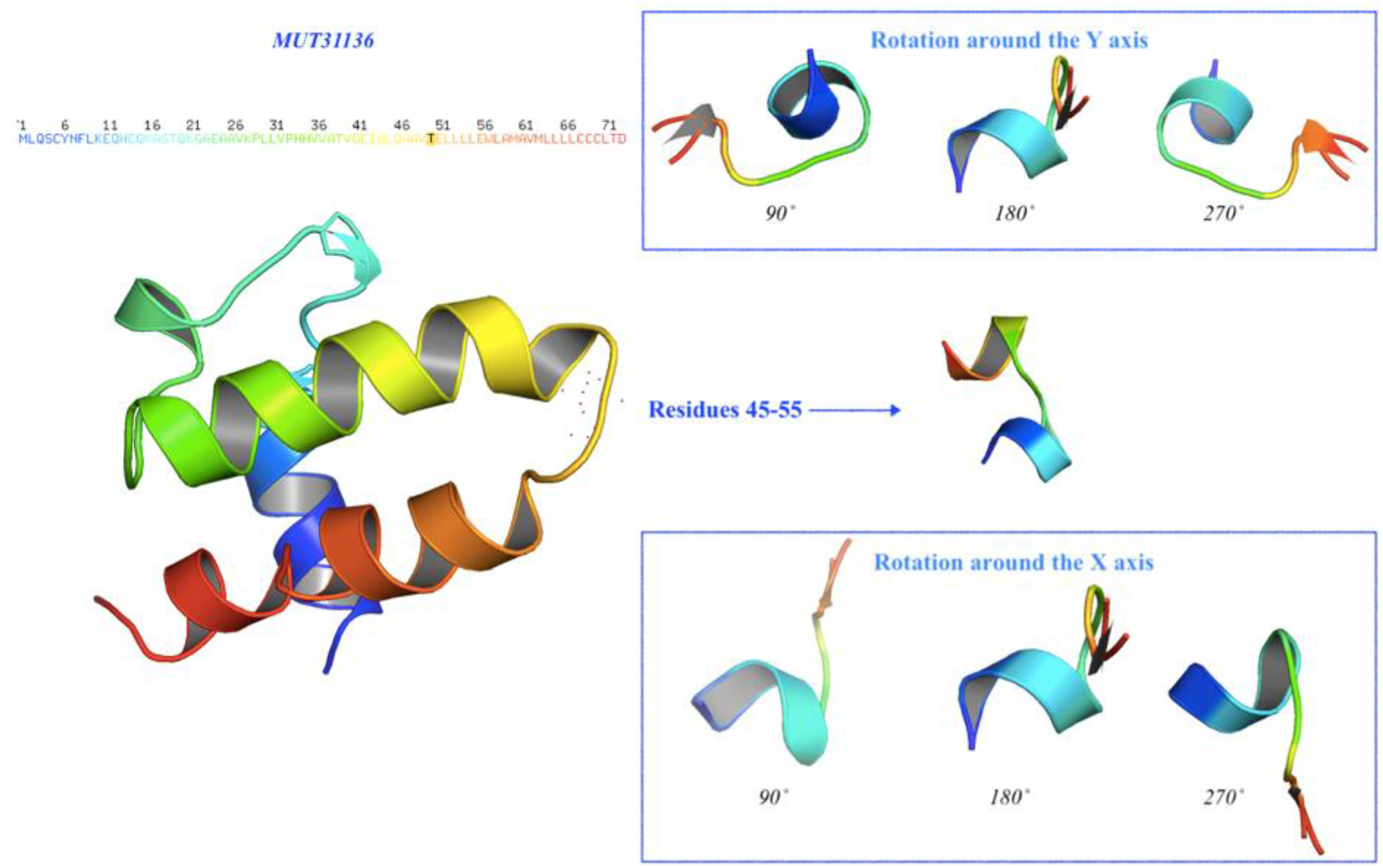
Prediction of the 3D structure for the mutated protein. The model MUT31136 represents the predicted tree-dimensional model of the protein subject to mutation. At the left side (top-down), the amino-acid sequence colored by the spectrum range, while in black color is the mutated amino-acid at position 50 (T). The protein has been oriented to facilitate the comparison and the residue 50 is represented with the red dots. At the right side (middle), detail of the residues 45-55 and their rotation around the Y axis (top) and around the X axis (bottom) with a step of 90°.

**Figure 8.**
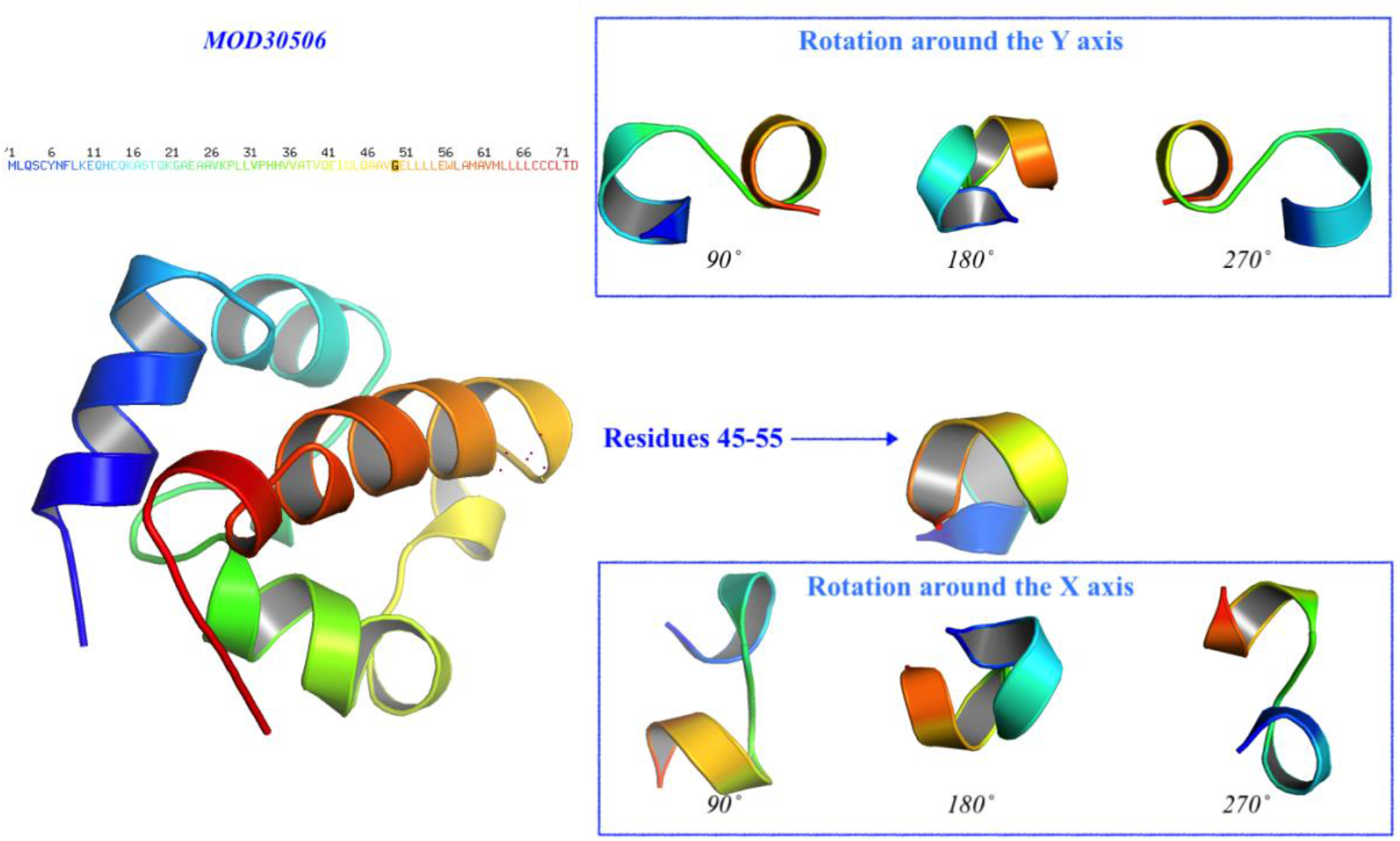
Prediction of the 3D structure for the unchanged protein. The model MOD30506 represents the predicted tree-dimensional model of the protein without mutation. At the left side (top-down), the amino-acid sequence colored by the spectrum range, while in black color is the investigated amino-acid at position 50 (G). The protein has been oriented to facilitate the comparison and the residue 50 is represented with the red dots. At the right side (middle), detail of the residues 45-55 and their rotation around the Y axis (top) and around the X axis (bottom) with a step of 90°.

### Protein related ontologies prediction

The analysis of the protein ontologies established different functions to the protein structure depending on whether it has mutated or not. As the Table 4 points out, the unchanged protein results related to catalytic and transferase activities with high probability (0.978 ≤ *p* ≤ 1). The mutated version of the protein presents a remarkable changing of its functionalities trend: even if usually the scores below 0.5 are interpreted as negative predictions, in an evolutionary context the decreasing of the probability of the transferase activity (from 0.98 to 0.375) to advantage of the binding function (from 0.004 to 0.132) sounds like not negligible. The contextual inversion of tendency of transferase to binding activity hints that the episodic evolutionary mutation aims to improve the binding ability.

**Table 4.** Classification report. The table displays the predicted functions of both (mutated and not) protein sequences and related scores. Only positive scores are reported, the ontological functions subject to inverted tendency are colored in red. “--” stays for unpredicted function.

| *GO terms* | Explanation | *Not mutated* | *Mutated* |
| --- | --- | --- | --- |
|  |  | Score | Score |
| *GO:0003674* | Molecular function | 1 | 1 |
| *GO:0003824* | Catalytic function | 1 | 0.998 |
| *GO:0016740* | Transferase activity | 0.978 | 0.375 |
| *GO:0016829* | Lyase activity | 0.017 | -- |
| *GO:0022891* | Transmembrane | 0.07 | -- |
| *GO:0005488* | Binding activity | 0.004 | 0.132 |
| *GO:0022892* | Transmembrane transport activity | 0.001 | 0.001 |

## Discussion and Conclusions

The recent worldwide outbreak of the COVID-19 disease is still ongoing and many researchers trying to make their input to the slowing down the spread of the disease as well as any other aspects such as study of epidemiology, pathogenesis, clinical presentation, treatment, and prevention. The main target for research is the SARS-CoV-2 virus, a novel coronavirus originated from Wuhan, China. To date, much is known about the structure, genetics and virulence of this virus, but many important questions are still not clear.

From the evolutionary analysis of SARS-CoV-2 now is known that the virus has the tendency to increasing mutation rate comparing to other *Coronaviridae*, such as SARS-COV, which has a moderate mutation rate(Zhao et al., 2004). The prevalence of single nucleotide transitions as the major mutational type across the world has been shown for SAR-CoV-2 virus (Mercatelli and Giorgi, 2020).

Our results shown that codon 9628 is under episodic selective pressure for all four countries. This leads that RNA secondary structure may be affected and consequently the protein product changes T (Threonine) to G (Glycine) in position 50 of the protein. This position is located close to the predicted transmembrane region. The mutation analysis revealed that change from G (Glycine) to D (Aspartic acid) may confer a new function to the protein, binding activity, which in turn may be responsible for attaching the virus to human eukaryotic cells. These can be accessed by *in vitro* experiments and possibly later facilitate a vaccine design and successful antiviral strategies.

Similar results presented by Mercatelli and Georgi have confirmed that clade G, prevalent in Europe, carries a D614G mutation in the Spike protein, which is responsible for the initial interaction of the virus with the host human cell. Other studies also showed different mutation location among different continents. Mutations in 2891, 3036, 14408, 23403 and 28881 positions are predominantly observed in Europe, whereas those located at positions 17746, 17857 and 18060 are exclusively present in North America (Pachetti et al., 2020). Their findings suggest that the virus is evolving and European, North American and Asian strains might coexist, each of them characterized by a different mutation pattern. Comparison of genomes from 13 countries also identified differences in protein coding sequences od SARS-CoV-2. For example, Indian strain showed mutation in spike glycoprotein at R408I and in replicase polyprotein at I671T, P2144S and A2798V, while the spike protein of Spain & South Korea carried F797C and S221W mutation, respectively (Khan et al., 2020). Moreover, the recently conducted integrative analyses of SARS-CoV-2 genomes from different geographical locations reveal unique features potentially consequential to host-virus interaction and pathogenesis (Sardar et al., 2020). However, the most recent to date study of genomic diversity and hotspot mutations in 30,983 SARS-CoV-2 genomes indicate that unlike the influenza virus or HIV viruses, SARS-CoV-2 has a low mutation rate which makes the development of an effective global vaccine very likely (Alouane et al., 2020). The study determined a number of hotspot mutations across the whole SARS-CoV-2 genome. Fourteen non-synonymous hotspot mutations (>10%) have been identified at different locations along the viral genome; eight in ORF1ab polyprotein (in nsp2, nsp3, transmembrane domain, RdRp, helicase, exonuclease, and endoribonuclease), three in nucleocapsid protein, and one in each of three proteins: Spike, ORF3a, and ORF8. Moreover, 36 non-synonymous mutations were identified in the receptor-binding domain (RBD) of the spike protein with a low prevalence (<1%) across all genomes (Alouane et al., 2020).

All these findings highlighted the importance of studying the relationship of geography of the isolates and mutations in their genomes because it is also can be confirmed by inferring from phylogenomic trees allowing to elucidate lineages and clusters accordingly to the geographical locations.

In conclusion, although we did not found correlation between countries located at the same meridian, we revealed that codon 9628 is under episodic selective pressure for all four countries.

## Acknowledgments

This work was supported by grants of Natural National Science Foundation of China (82020108022, Shu-Lin Liu). The funding bodies played no roles in the design of the study; the collection, analysis or interpretation of data; or in writing the manuscript.

## Declarations of interest

none

## Notes

### Competing Interest Statement

The authors have declared no competing interest.

